# Charting human subcortical maturation across the adult lifespan with *in vivo* 7 T MRI

**DOI:** 10.1101/2021.04.07.438765

**Authors:** Steven Miletić, Pierre-Louis Bazin, Scott J. S. Isherwood, Max C. Keuken, Anneke Alkemade, Birte U. Forstmann

## Abstract

The human subcortex comprises hundreds of unique structures. Subcortical functioning is crucial for behavior, and disrupted function is observed in common neurodegenerative diseases. Despite their importance, human subcortical structures continue to be difficult to study *in vivo*. Here we provide a detailed account of 17 prominent subcortical structures, describing their approximate iron and myelin contents, morphometry, and their age-related changes across the normal adult lifespan. The results provide compelling insights into the heterogeneity and intricate age-related alterations of these structures. They also show that the locations of many structures shift across the lifespan, which is of direct relevance for the use of standard magnetic resonance imaging atlases. The results further our understanding of subcortical morphometry and neuroimaging properties, and of normal aging processes which ultimately can improve our understanding of neurodegeneration.

## 1 Introduction

The human subcortex comprises hundreds of unique structures (Alkemade, Keuken, & Forstmann, 2013; Forstmann, de Hollander, Van Maanen, Alkemade, & Keuken, 2017) which receive interest from a broad range of neuroscientific disciplines (e.g. Lozano et al., 2019; Raznahan et al., 2014; G. M. Shepherd, 2013; Tian, Margulies, Breakspear, & Zalesky, 2020). Subcortical functioning is crucial for normal behavior and physiology including decision making (Ding & Gold, 2013), reward processing (O’Doherty et al., 2004; Schultz, Dayan, & Montague, 1997), and motor behavior (Mink, 1996). Disruption of subcortical structures is observed in common neurodegenerative diseases including Parkinson’s (Hirsch, Graybiel, & Agid, 1988) and Alzheimer’s disease (Ehrenberg et al., 2017; German, White, & Sparkman, 1987). Subcortical structures are also of interest as (potential) deep brain stimulation (DBS) targets in Parkinson’s disease (Fasano & Lozano, 2015; Limousin et al., 1995) and other disorders such as major depression and epilepsy (Lozano et al., 2019).

Research into the subcortex depends on the imaging of individual subcortical structures. However, visualizing subcortical structures using *in vivo* methods such as magnetic resonance imaging (MRI) is challenging due to their close spatial proximity, biophysical properties, and morphometry (Keuken, Isaacs, Trampel, van der Zwaag, & Forstmann, 2018). As a consequence, our understanding of the subcortex remains limited, and lags behind our understanding of the cortex. Quantitative ultra-high field 7 Tesla MRI provides a method to overcome the challenges associated with visualizing subcortical structures (Bazin, Alkemade, Mulder, Henry, & Forstmann, 2020; Keuken et al., 2018), which we use here to provide a cross-sectional account of the subcortex across the adult lifespan.

The biophysical properties that determine the appearance of brain structures on MR images include the iron and myelin contents, which influence t he m ain s ources of contrast in MRI: the longitudinal and effective transverse relaxation rates, and the local susceptibility to magnetic fields. Furthermore, iron and myelin are highly biologically relevant: Myelin plays an important role in plasticity and development (e.g. Fields, 2015; Hill, Li, & Grutzendler, 2018; Turner, 2019), and iron is crucial for normal tissue functioning (e.g. Zecca, Youdim, Riederer, Connor, & Crichton, 2004). Iron deposition (Daugherty & Raz, 2013; Hallgren & Sourander, 1958; Raz & Rodrigue, 2006; Ward, Zucca, Duyn, Crichton, & Zecca, 2014; Zecca et al., 2004) and decreased myelination (Raz & Rodrigue, 2006; Shen et al., 2008) are part of normal aging processes, but excessive iron accumulation and myelin degradation are prominent in diseases including Parkinson’s and Alzheimer’s disease (e.g. Mancini et al., 2020; Zecca et al., 2004). A description of normal age-related changes in iron and myelin content can therefore provide a frame of reference to contrast pathological iron accumulation and myelin degradation, and to refine methods for the early detection of pathological alterations using MRI measures as biomarkers.

An additional factor determining the appearance of the human subcortex is the small size of the individual structures. Prominent subcortical structures such as the subthalamic nucleus are as small as a few millimeters thick, limiting the number of voxels they encompass on MR images commonly used in research and in the clinic. Moreover, voxels at the border of structures likely include tissue from adjacent structures (*partial voluming*), which can lead to biases especially when voxel sizes are large relative to the structure (Mulder, Keuken, Bazin, Alkemade, & Forstmann, 2019). Structure size should therefore be taken into account when imaging the subcortex. An important additional consideration here is the development of atrophy with increasing age, which is reflected in reduced volume of grey matter structures (Cherubini, Péran, Caltagirone, Sabatini, & Spalletta, 2009; Courchesne et al., 2000; Herting et al., 2018; Lemaitre et al., 2012; Raz, 2004; Raz & Rodrigue, 2006; Walhovd et al., 2005) and which results in more cerebrospinal fluid (CSF) and larger ventricles (Good et al., 2001; Greenberg et al., 2008; Stafford, Albert, Naeser, Sandor, & Garvey, 1988; Walhovd et al., 2005). In addition to volume changes, atrophy can result in a shift in the location of structures (Keuken et al., 2013; Keuken et al., 2017; Kitajima et al., 2008).

These factors combined hamper visualization of the subcortex when using conventional magnetic resonance imaging techniques. Furthermore, the age-related alterations in these factors alter the appearance of the subcortex with increasing age. In this study, we provide a detailed account of 17 subcortical structures using data from 105 healthy participants across the adult lifespan obtained with *in vivo* methods tailored for studying the human subcortex (Alkemade, Mulder, et al., 2020). We first estimate the iron and myelin distributions and size of individual structures. We next analyze age-related changes: iron deposition, myelin degradation, and the morphometric changes that are the result of atrophy (changes in size, shape, and location).

## 2 Results

One hundred and five healthy volunteers were scanned using an ultra-high field 7 Tesla MRI scanner as part of the openly available Amsterdam ultra-high field adult lifespan database project (AHEAD; Alkemade, Mulder, et al., 2020). A quantitative, multi-modal MP2RAGE-ME sequence (Caan et al., 2019) with 0.7 mm isotropic resolution was used to simultaneously estimate R1, R2* and quantified susceptibility mapping (QSM) values in a single scanning sequence. For each participant, 17 subcortical structures (see Table 1) were delineated using the Multi-contrast Anatomical Subcortical Structures Parcellation method (MASSP; Bazin et al., 2020).

**Table 1:**
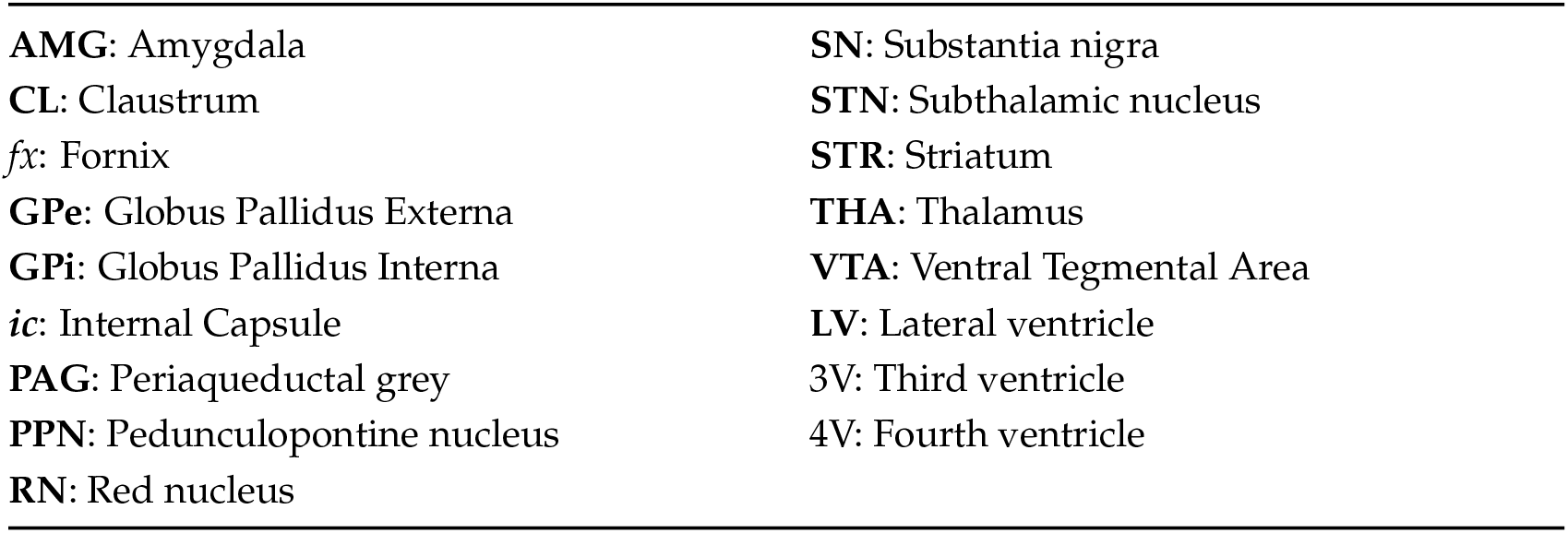
Regions of interest. Midline structures were parcellated as a single structure, all other structures (indicated by bold-faced letters) were parcellated separately per hemisphere. Abbreviations in italics indicate white matter structures.

We analyzed each structure by first estimating the *iron* and *myelin* concentrations, using simplified biophysical models that translate the measured R 1, R 2*, and QSM values into the most likely corresponding iron and myelin concentrations (see Methods). We obtained both the (within-structure) median of iron and myelin distributions, and the interquartile range (IQR) which reflects image noise and tissue (in)homogeneity. Second, we analyzed the structure morphometry by estimating *volume* and *thickness*. Thickness is defined as twice the distance between the boundary and the internal skeleton of the structure. As a local measure (contrary to volume), it is defined for every voxel in a structure, and it depends on the structure’s shape. Also for thickness, we determined both the median and IQR, the latter reflecting the regularity of the structure’s shape: Regularly shaped structures (e.g., the red nucleus) have a similar thickness at each voxel’s location, resulting in lower between-voxel IQRs compared to complex shaped structures (e.g., the striatum). Third, we determined the *location* (center of mass in 3 Cartesian coordinates) of each structure. Center of mass was determined after applying an affine transformation to a group template, to account for inter-individual differences in intracranial volume and shape, while retaining inter-individual variability in distances relative to the neurocranium.

The distributions of iron, myelin, and volumes revealed a large between-structure heterogeneity in the human subcortex (Figure 1). The globus pallidus externa and interna, red nucleus, substantia nigra, and subthalamic nucleus displayed the highest iron concentrations (both median and IQR), corroborating earlier reports (Haacke et al., 2005; Hallgren & Sourander, 1958; Ramos et al., 2014). In line with expectations, low iron concentrations in combination with high myelin concentrations were observed in the white matter structures under study: the internal capsule and the fornix. The estimated myelin concentrations of the subthalamic nucleus, red nucleus, and ventral tegmental area were relatively high, which causes the limited visibility of these structures on T1-weighted images (Keuken et al., 2018). For comparison, the estimated myelin concentrations of the striatum and amygdala were substantially lower, resulting in intensities comparable to cortical grey matter on T1-weighted images. The within-structure IQR of iron roughly scaled with the median estimates, indicating that structures with high iron concentrations tended to be less homogeneous. This also held for the myelin concentrations, with the exception of the fornix, which had a particularly high IQR. This could potentially have been caused by partial voluming with the lateral ventricles, decreasing the myelin estimates at voxels near the boundary of the fornix. Finally, the striatum, thalamus, lateral ventricles, and internal capsule were the largest structures, and the subthalamic nucleus the smallest.

**Figure 1:**
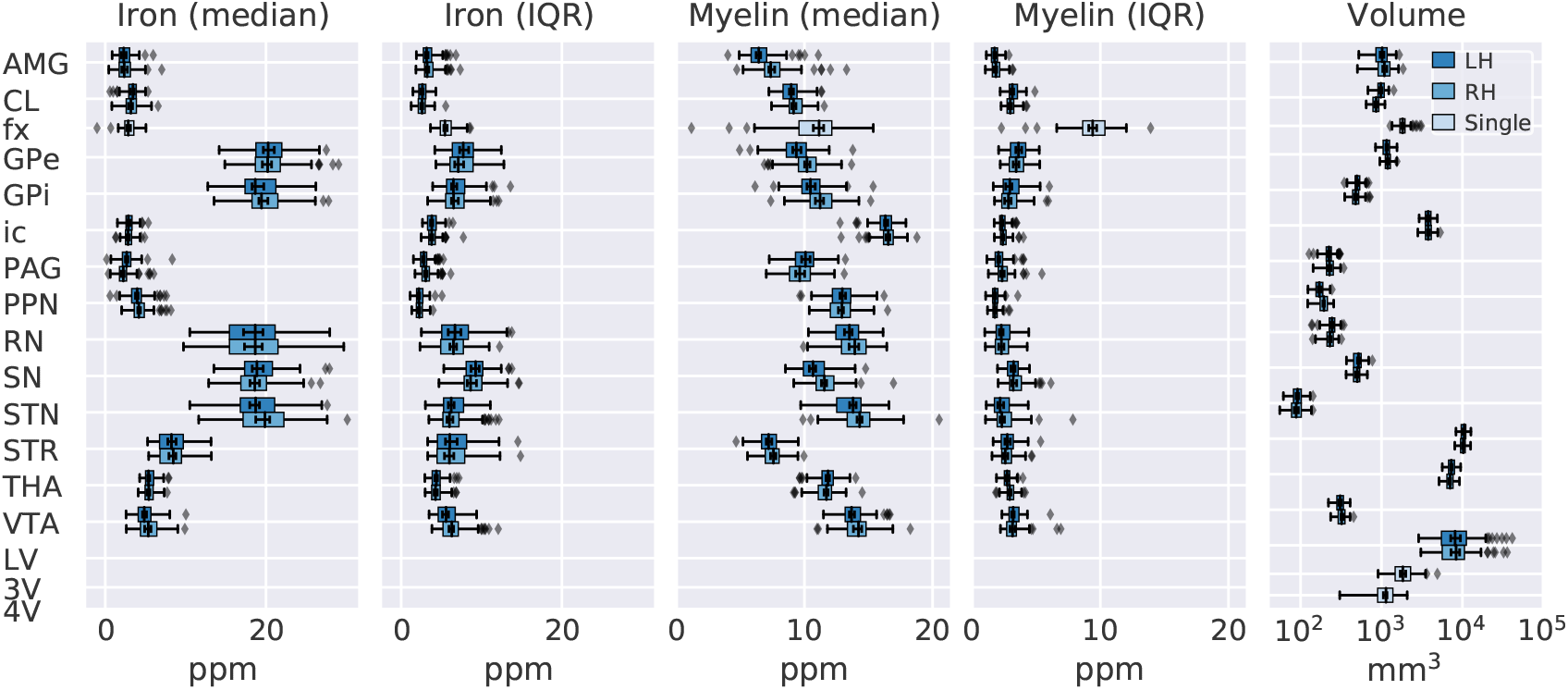
Across-participant distributions of within-structure median and IQR of iron and myelin, and volumes per structure. The center line in each box marks the median, box limits are the across-participant IQR, and whiskers are at 1.5 times the IQR below and above the box limits. Error bars drawn inside boxes indicate 95% confidence intervals around the median, obtained by bootstrapping with 10,000 iterations. Colors indicate hemisphere (LH = left hemisphere, RH = right hemisphere, Single = structures that are continuous across the hemispheres), ppm = parts per million, mm = millimeter.

### 2.1 Maturation effects

We next studied the age-related alterations in iron, myelin, and morphometry across the adult lifespan. We fit a set of 24 regression model specifications (with, as predictor variables, linear and/or quadratic effects of age, plus sex and potential interactions between sex and age) for all structures and measures individually. As we had no *a priori* hypothesis on lateralization, we collapsed across hemispheres to reduce the total number of fitted models. The model specification that showed the most parsimonious trade-off between quality of fit and model complexity (as quantified using the Bayesian information criterion; Schwarz, 1978) was considered the winning model and used for further analyses. To help navigate the winning models of each structure and measure (including R1, R2*, and QSM values), we developed an online interactive app, which is accessible at https://subcortex.eu/app (see also Figure S1).

We observed (median) iron accumulation in all structures except for the claustrum, globus pallidus interna, and periaqueductal gray, which instead showed stable iron concentrations (Figure 2). With the exception of the globus pallidus interna, the iron-rich basal ganglia appeared to accumulate most iron during aging in absolute terms. The IQRs increased with age for all structures, revealing a global decrease in structure homogeneity. Since this decrease in homogeneity was also present in the structures where no median iron increase was observed, it likely partially reflects an increase in image noise. However, the increases in IQR were substantially higher in the structures that accumulate most iron, suggesting that the (median) iron accumulation for these structures was not homogeneously distributed within the structure. This decrease in homogeneity was particularly strong in striatum and the red nucleus.

**Figure 2:**
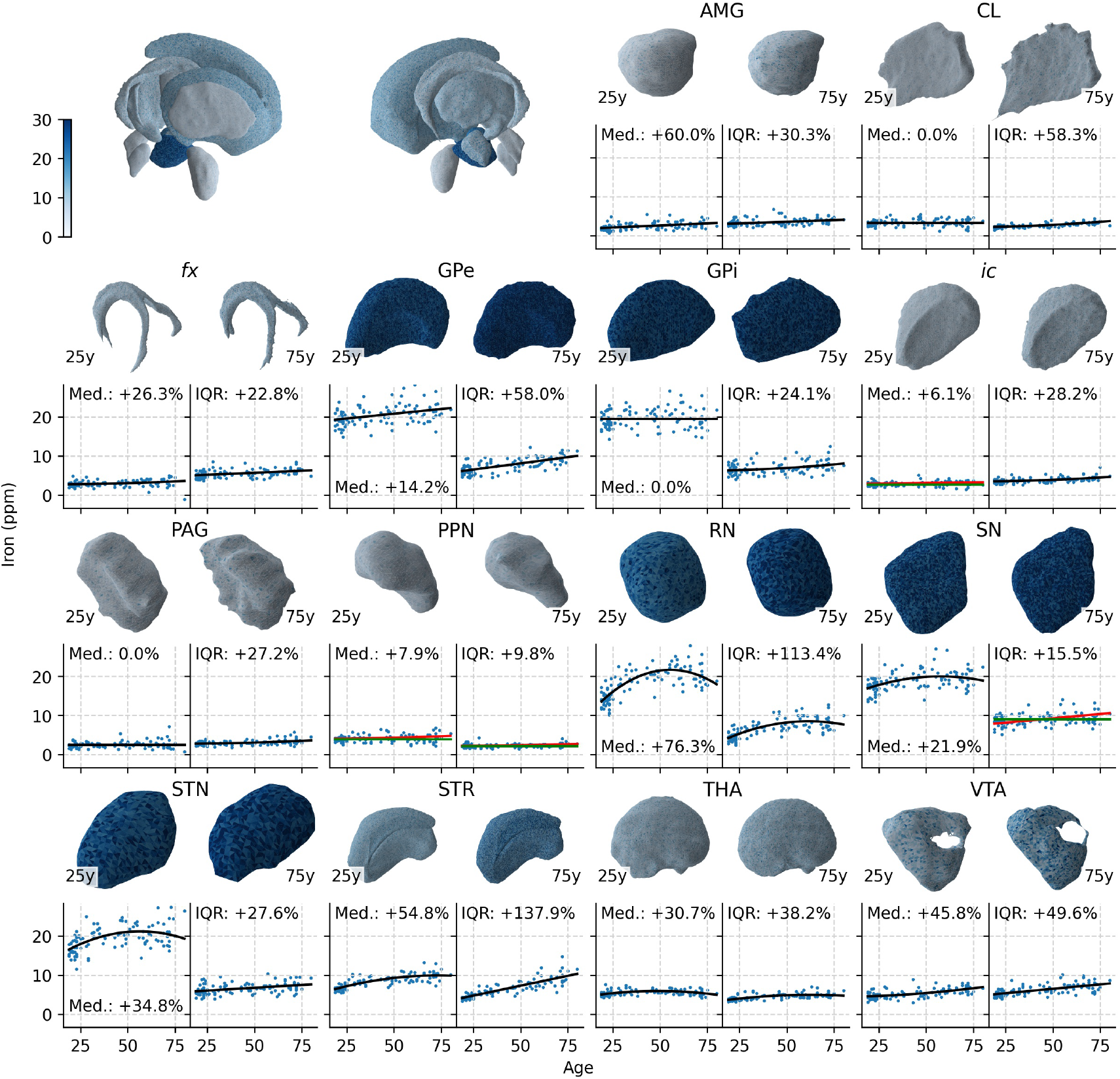
Age-related changes in iron content. The meshes are based on the young (18–30 years old; left) and elderly (70–80 years old; right) participants after a non-linear transformation to MNI2009b space. Mesh colors illustrate the model predictions for the median and IQR of iron distributions at 25 (left) and 75 (right) years old. Colors in the top-left meshes of all structures indicate model predictions at 25 years old. In case the winning model did not include sex as a predictor variable, the model predictions are shown in black lines; otherwise, green and orange lines are used for the predictions for women and men, respectively. The total amount of change in median (Med.) and IQR are shown in each scatterplot. The ventricles are assumed to have no iron and are excluded from this graph.

In line with expectations, we observed a general myelin degradation (see Figure 3), except for in the amygdala, claustrum, and substantia nigra, where no alterations in myelin concentrations were detected. The largest (absolute) reduction of myelin was present in the fornix; the other white matter structure, internal capsule, showed a smaller decrease in myelin. The globus pallidus interna, periaqueductal grey, pedunculopontine nucleus, substantia nigra, and ventral tegmental area showed slightly higher median myelin concentrations in females than in males. Like in the case of iron, the increases in IQR of myelin reflected a trend of decreasing structure homogeneity across structures. Since these IQR increases were present for structures that did not show any change in median myelin content, they likely partially reflect increases in image noise. However, the structures with large median myelin decreases also tended to show the largest increases in IQR, suggesting that these IQR increases also reflect increasing inhomogeneities in myelin concentrations.

**Figure 3:**
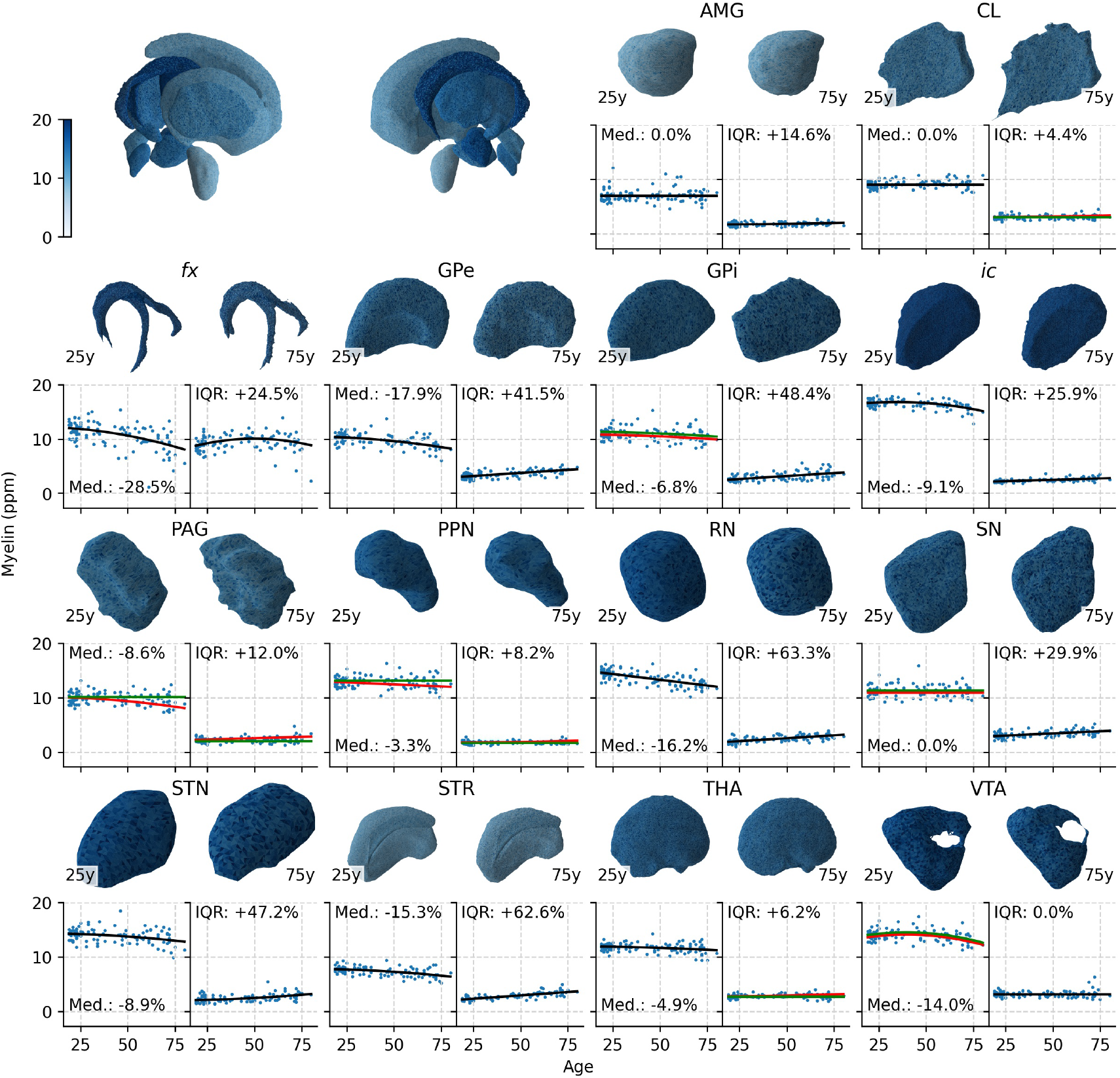
Age-related changes in myelin content. The meshes are based on the young (18–30 years old; left) and elderly (70–80 years old; right) participants after a non-linear transformation to MNI2009b space. Mesh colors illustrate the model predictions for the median and IQR of myelin distributions at 25 (left) and 75 (right) years old. Colors in the top-left meshes of all structures indicate model predictions at 25 years old. In case the winning model did not include sex as a predictor variable, the model predictions are shown in black lines; otherwise, green and orange lines are used for the predictions for women and men, respectively. The total amount of change in median (Med.) and IQR are shown in each scatterplot. The ventricles are assumed to have no myelin and are excluded from this graph.

Next, we analyzed the effects of atrophy (Figure 4). The lateral and third ventricle showed a substantial volume increase with age, which can at least partially be explained by the filling of the intracranial space created by a trophied brain tissue. Contrary to expectations, the volume of the fourth ventricle decreased rather than increased. Inspection of the mesh of the fourth ventricle in the elderly suggests this may be caused by shrinkage of the superior part. Volume decreases were also found in the striatum, thalamus, amygdala, ventral tegmental area, periaqueductal grey, pedunculopontine nucleus, and red nucleus, likely reflecting a trophy. The internal capsule, fornix and globus pallidus interna showed a small increase in volume with age, suggesting white matter swelling, which could be caused by neuroinflammatory processes.

**Figure 4:**
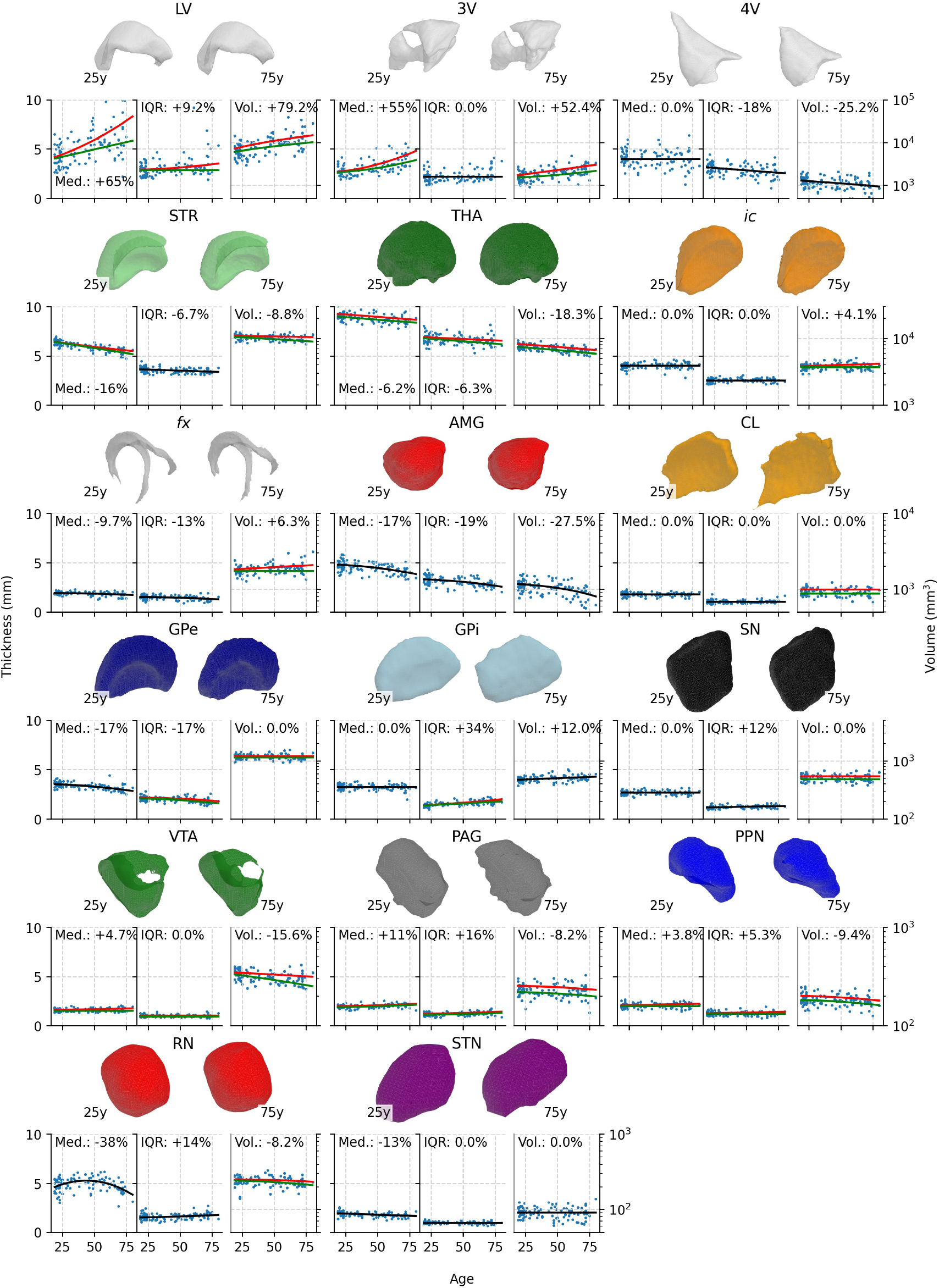
Age-related changes in structure morphometry. The meshes are based on the young (18–30 years old; left) and elderly (70–80 years old; right) participants after a non-linear transformation to MNI2009b space. The lines in each scatterplot visualize the winning model predictions. In case the winning model did not include sex as a predictor variable, the model predictions are shown in black lines; otherwise, green and orange lines are used for the predictions for women and men, respectively. The total amount of change in median (Med.) and IQR of thickness and volume (Vol.) are shown in each scatterplot.

Atrophy of specific subparts of a structure, as a result of increased vulnerability to atrophy in that part, could result in shape changes (Ho et al., 2020; Raznahan et al., 2014). Shape changes can be detected by analyzing changes the median and IQR of thickness, which depend on the structure’s shape. Specifically, when changes in the median thickness and volume point in the same direction (as is the case in, e.g., the lateral ventricles, striatum), this suggests overall thickening or thinning of a structure. Instead, increases in median thickness combined with decreases in volume can indicate atrophy in a thinner part of the structure, as this would decrease the amount of voxels with relatively low thickness, increasing the median thickness. This specific effect appeared to be present in the ventral tegmental area, pedunculopontine nucleus and periaqueductal grey. Furthermore, increases in IQR indicate decreases in structure regularity, which was observed in the globus pallidus interna, substantia nigra, periaqueductal grey, pedunculopontine nucleus, and red nucleus.

A third potential effect of atrophy is a change in the location of individual structures relative to the neurocranium (Keuken et al., 2013; Keuken et al., 2017; Kitajima et al., 2008): As the brain atrophies, the resulting physical space is filled with CSF, leading to location shifts of other brain structures. For the majority of brain structures under investigation, we observed location shifts in the lateral and inferior direction (Figure 5). The center of mass of the lateral and third ventricles and the claustrum also shifted in the posterior direction; the fornix and striatum shifted in anterior direction.

**Figure 5:**
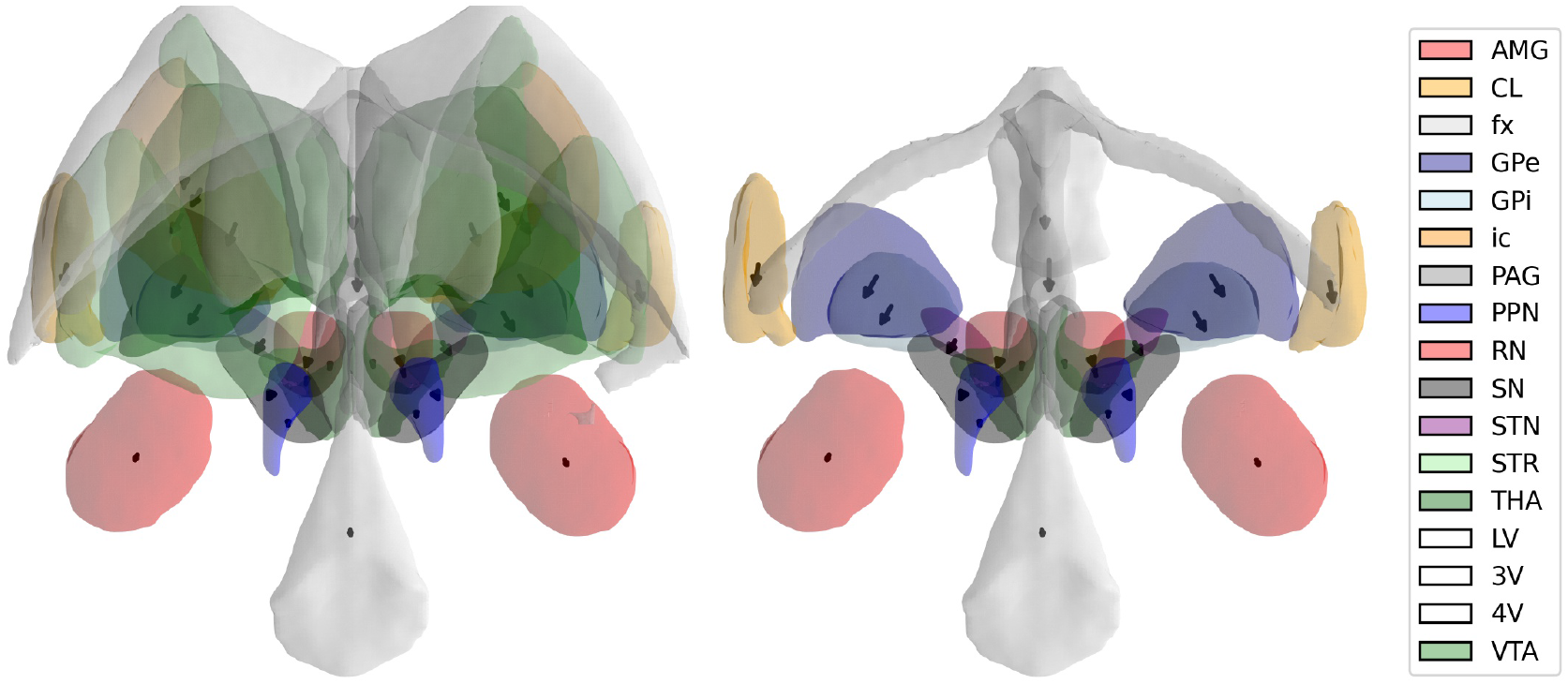
Age-related changes in structure location, posterior view. Meshes were based on the young (18–30 years old) participants after non-linearly transforming to MNI2009b space. Arrows depict the model predictions for the location shift, starting at the center of mass of each structure in MNI2009b space, and pointing to the predicted center of mass of the structure at 75 years old. The left graph shows all 17 subcortical structures under investigation, the right graph excludes the lateral ventricles, internal capsule, and thalamus, to improve the visibility of the smaller structures.

Combined, we observed age-related changes in all measures: iron, myelin, volume, thickness, and location. These effects were in line with the expected effects of iron accumulation, myelin degradation, and atrophy, but there appeared to be strong between-region variability in the degree to which regions change with age, which we focus on in the next section.

### 2.2 Between-structure variability in maturation

Because the winning models of age-related change trajectories included either linear or quadratic influences of age, the parameter estimates of the different models cannot be directly compared. To provide a quantity that summarizes the amount of age-related change (irrespective of the underlying model specification), we calculated the sum of the absolute yearly changes between 19 and 75 years old, relative to the model’s predicted value at 19 years old to take into account baseline differences. This *total age-related change* metric (also shown in Figures 2–4 and S2) revealed that the strongest age-related changes, across metrics, were observed in the striatum and red nucleus, driven by tissue degeneration and iron accumulation (Figure 6). The pedunculopontine nucleus, claustrum, and internal capsule remained relatively stable across the age range.

**Figure 6:**
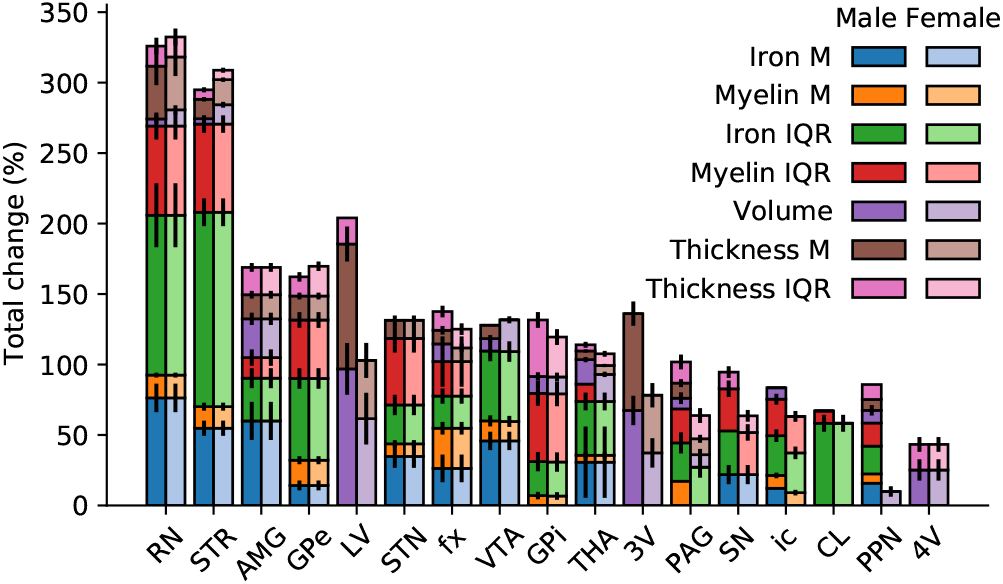
Total change per morphometric measure between 19 and 75 years, relative to the measure’s predicted value at 19. Error bars indicate standard errors, obtained by bootstrapping with 10,000 iterations.

Sex differences in aging effects were present in the measures that reflect atrophy, where we observed larger ventricular volume and thickness increases in males compared to females, and slightly larger volume and thickness decreases in striatum, ventral tegmental area, and red nucleus in females compared to males. Sex differences in atrophy rates have been observed before, although the size and direction of sex effects appears to depend on the specific structure (Li et al., 2014; Wang, Xu, Luo, Hu, & Zuo, 2019; Xu et al., 2000). In the iron and myelin measures, sex differences were relatively modest and confined to specific regions. The most prominent were in the thalamus (males but not females showed a decrease in the IQR of myelin), the periaqueductal grey (males but not females showed median myelin decrease and IQR increase), internal capsule (males but not females showed median myelin decrease), and pedunculopontine nucleus (males but not females showed alterations in iron and myelin). These sex differences, however, were modest, and the overall aging patterns were comparable across sexes.

Finally, we tested whether groups of structures with correlated aging patterns could be identified. We computed all pairwise correlations between structures using their total age-related changes (in median and IQR in iron, myelin, and thickness, as well as volume) as outcome variables (i.e., using the matrix visualized in Figure S2 as input, excluding the location changes). The correlation matrix was then organized using data-driven hierarchical clustering, the result of which is shown in Figure 7. The predominantly positive correlations reiterate the presence of a global, across-region aging process. Furthermore, there are clusters of regions that covary strongly. One such cluster consists of the ventricular system, which ages differently compared to other regions, due to the volume increases rather than decreases. Contrary to expectations, the other clusters that were identified did not group anatomically or functionally related regions such as the basal ganglia or the dopaminergic midbrain structures, although some functionally related nodes do show high pairwise correlations (e.g., subthalamic nucleus and substantia nigra). Instead, the largest cluster of regions with similar aging patterns included the striatum, red nucleus, claustrum, internal capsule, globus pallidus externa.

**Figure 7:**
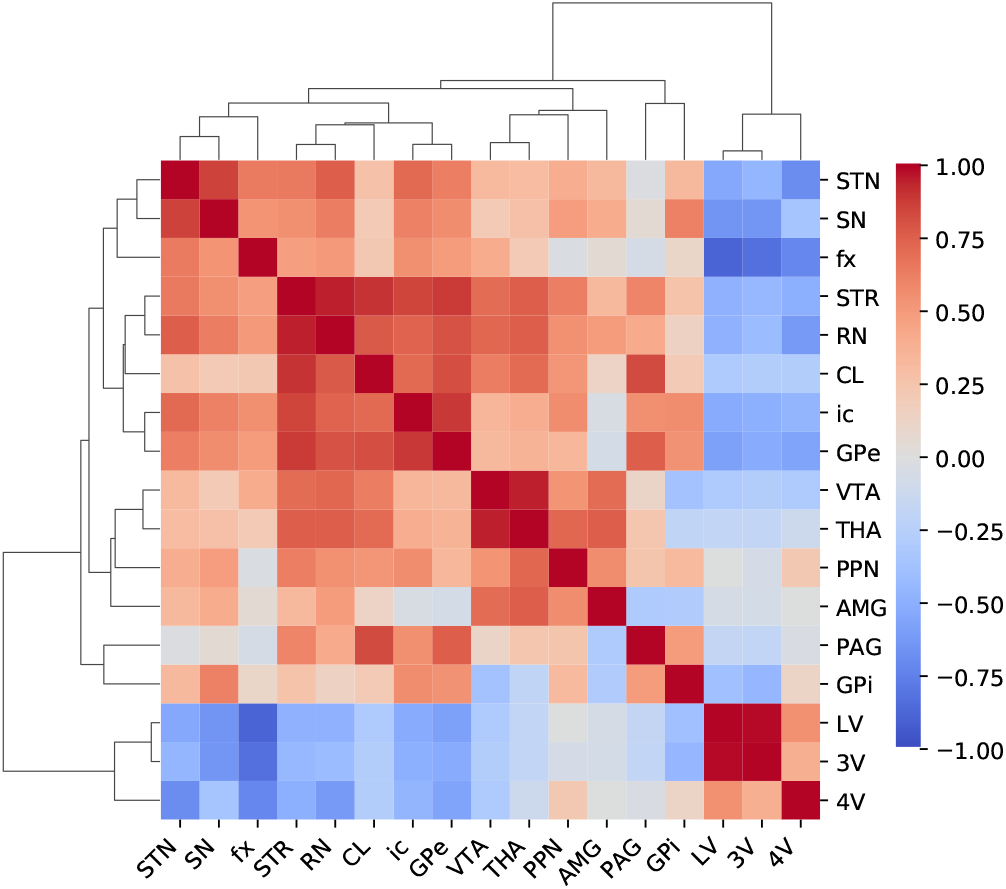
Correlation matrix indicating the similarities and differences between structures’ aging patterns. Regions are ordered using hierarchical clustering, of which the dendrograms are shown on the left and top of the correlation matrix. Colors indicate correlation coefficients.

## 3 Discussion

Interest in the human subcortex is rapidly growing in cognitive and clinical neuroscience due to the relevance of subcortical regions as (potential) targets for DBS and their role in cognition. Here, we studied 17 subcortical structures in terms of their iron and myelin contents, their sizes, as well as the intricate age-related alterations. Our results highlight the heterogeneity in the subcortex, presenting the strong variability in iron, myelin, and morphometry that exists between structures. Furthermore, our results indicate global effects of iron accumulation, myelin degradation, and atrophy in the subcortex across the normal adult lifespan, and strong variability in the extent to which the different structures are affected by each type of age-related change. The red nucleus and striatum show most profound changes, whereas some other structures (such as the claustrum and pedunculopontine nucleus) appear less affected by aging processes.

Furthermore, we observe clusters of regions with similar aging patterns, but these clusters (except for the ventricular system) do not show similarity between the aging patterns of groups of regions that are functionally or anatomically related. For example, it could be hypothesized that there are similarities between the aging processes of the globus pallidus externa and subthalamic nucleus, based on their structural connectivity, or between the ventral tegmental area and substantia nigra, as these are the dopaminergic midbrain areas. However, the correlations in aging patterns between such related regions were not particularly strong, and substantially weaker than those between regions that are anatomically and functionally less related such as the thalamus and the ventral tegmental area. This suggests that functional or anatomical relations between regions do not imply similar aging patterns, and that inferences about aging of one region cannot be drawn on the basis of aging patterns of similar regions.

To better navigate the rich landscape of subcortical aging, we also share our results in an online app (Figure S1, https://subcortex.eu/app) that can be used to create interactive and intuitive 3D visualizations of the human subcortex across the lifespan and across modalities. It allows for inspection and reuse of the underlying models and data of each individual structure. The app was designed in a flexible way, so that it can be augmented with more structures and contrasts to expand it to a comprehensive chart of the human subcortex. The underlying data can readily be downloaded for further analyses.

Understanding the aging processes in the subcortex is paramount in research and in clinical practice. While iron accumulation and myelin degradation are part of normal aging processes, increased accumulation and myelin degradation are part of multiple neurodegenerative disorders including Parkinson’s and Huntington’s disease (Andersen, Johnsen, & Moos, 2014; Collingwood & Davidson, 2014; Ward et al., 2014; Zecca et al., 2004). An accurate description of the distributions of iron and myelin across the lifespan in health provides a frame of reference against which pathological iron accumulation and myelin degradation can be contrasted, and can prove useful in the development of biomarkers for disease (Guan, Xu, & Zhang, 2017; Mancini et al., 2020; Martin, Wieler, & Gee, 2008; Schenck & Zimmerman, 2004; Zecca et al., 2004).

Iron and myelin are also the two main determinants of MRI contrast. Many subcortical structures, such as the subthalamic nucleus, cannot readily be distinguished on conventional T1-weighted MRI images due to a lack of contrast with nearby regions. Because of the limited visibility of subcortical structures on conventional MR images, a common practice is to use atlases to locate individual structures (Devlin & Poldrack, 2007; Evans, Janke, Collins, & Baillet, 2012). Stereotactic atlases based on *post mortem* tissue are often used for planning DBS surgery, and probabilistic MRI atlases are conventionally used in cognitive neuroscientific research. Subcortical MRI atlases are growing in numbers (Keuken et al., 2014; Lau et al., 2020; Pauli, Nili, & Tyszka, 2018; Trutti et al., 2021; Ye et al., 2021) due to improvements in MRI resolution and contrasts. However, MRI atlases are typically developed using anatomical images obtained from young participants, which can cause biases when such atlases are subsequently used to infer anatomical information in older participants or patient populations (Evans et al., 2012; Keuken et al., 2013; Samanez-Larkin & D’Esposito, 2008). In cognitive neuroscience research, it is common to apply spatial normalization procedures to a group space to account for individual differences in anatomy, but consistent deviations from the group template are likely to introduce normalization errors (Samanez-Larkin & D’Esposito, 2008). These biases can result from iron accumulation and myelin degradation (which change the contrast of images) and from atrophy (which change the size and the location of structures). Our results can help understand the biases that could occur when conventional MRI atlases, based on young participants, are used to analyze data from older participants, and call for the development of age-specific MRI atlases of the subcortex to remedy these biases.

The between-region variation in iron contents has important consequences for blood oxygenation level dependent (BOLD) functional MRI. Because iron decreases T2* relaxation times, on which contrast-to-noise ratios (CNR) of BOLD-fMRI sequences depend, BOLD CNR varies substantially between regions. For instance, within young participants, the CNR in the red nucleus is expected to be 72% lower than in the amygdala, when using an echo time of 42 ms at 7 T (corresponding to the T2* of the amygdala in young participants), solely due to the differences in T2* (see supplement for details). Age-related alterations in iron contents can have similar effects. For instance, the CNR in the red nucleus at 50 years old is 28% lower than at 19 years old when using an echo time of 18 ms at 7 T (corresponding to the T2* of the red nucleus at 19 years old). Thus, iron deposition can confound fMRI studies into age-related changes of BOLD responses.

However, substantial gains in CNR can be achieved by optimizing the echo time to meet the specific requirements of studying a structure of interest (see also de Hollander, Keuken, van der Zwaag, Forstmann, & Trampel, 2017; Miletić, Bazin, et al., 2020). For instance, when targeting the red nucleus, decreasing the echo time to 18 ms (corresponding to the T2* of the red nucleus at 19 years old) is expected to lead to a 58% higher CNR compared to an echo time of 42 ms (which would be optimal to target the amygdala). Similarly, the echo time can be adjusted to partially mitigate the effects of age-related changes in T2*: By decreasing the echo time from 18 ms to 13.7 ms (corresponding to the T2* of the red nucleus at 50 years old), a modest 5% increase in CNR can be expected. Using our online app as a resource for participant-specific predictions of R1, R2*, and QSM values, we envision the use of MRI protocols tailored to the structure of interest and the participant’s age and sex.

The present study has several limitations. First, iron and myelin contents were approximated using simplified biophysical models (c.f. Hametner et al., 2018; Mangeat, Govindarajan, Mainero, & Cohen-Adad, 2015; Marques, Khabipova, & Gruetter, 2017; Metere & Möller, 2018; Rooney et al., 2007; Stüber et al., 2014), which rely on population-average iron and myelin estimates found in the literature (Hallgren & Sourander, 1958; Metere & Möller, 2018; Randall, 1938). Since the amount of structures for which myelin concentrations are known is limited, we also relied on *post mortem* materials to infer myelin concentrations. While the biophysical models provided a good fit to our data, the resulting iron and myelin estimates could be biased by the relatively limited information on myelin concentrations in the literature. Such potential biases should be present for all participants and therefore not affect the age-related alterations we report. The approach of estimating myelin and iron concentrations based on quantitative MRI using simplified biophysical models would benefit from more quantitative myelin and iron estimates obtained through analyzing *post mortem* tissue.

Second, we cannot exclude age-related changes in parcellation accuracy. We relied the MASSP algorithm (Bazin et al., 2020) to parcellate the 17 subcortical regions in each participant individually. The performance of MASSP (compared to manual delineations) tends to decrease with age. Fortunately, the impact of age-related biases in parcellation was shown to be limited for the quantitative MRI measures (Bazin et al., 2020) on which the iron and myelin estimates are based, suggesting that the age-related changes in myelin and iron are unlikely to be caused by age-related differences in parcellation performance. On the other hand, size estimates (volume and to a lesser extent thickness) and more susceptible to the age-related changes in parcellation quality. Here, we used an improved version of MASSP to mitigate these effects, although we cannot exclude any remaining age-related decreases in parcellation accuracy.

Our results extend previous studies into aging patterns of the subcortex, which focus on a smaller number of typically large subcortical areas, often based on MRI with lower field strengths (Aquino et al., 2009; Cherubini et al., 2009; Daugherty & Raz, 2013, 2016; Fjell et al., 2013; Greenberg et al., 2008; Herting et al., 2018; Keuken et al., 2017; Li et al., 2014; Raz, 2004; Raz et al., 2005; Raz & Rodrigue, 2006; Raznahan et al., 2014; Walhovd et al., 2005; Wang et al., 2019). Experiments using very large numbers of participants detected complex nonlinear age-related changes in some subcortical structures (Fjell et al., 2013; Raznahan et al., 2014), suggesting the presence of critical ages that demark changes in the rates of atrophy in some subcortical areas (Fjell et al., 2013). Our study had a more modest sample size, which did not allow to evaluate complex non-linear trends. On the other hand, by leveraging an open database of ultra-high field 7 T quantitative MRI, we could provide a first view on many structures and variables at once, which may be refined as more 7 T quantitative MRI data sets become available. As such, our study provides a richer and more extensible description of subcortical composition, morphometry and aging.

## 4 Methods

### 4.1 Participants

We used the Amsterdam ultra-high field adult lifespan database (AHEAD; Alkemade, Mulder, et al., 2020), which consists of multimodal MRI data from 105 healthy participants. Inclusion criteria were age 18–80 years and self-reported health at the time of inclusion. Exclusion criteria were any factors that could potentially interfere with MRI scanning, including MRI incompatibility (e.g., pacemakers), pregnancy, and self-reported claustrophobia. At least six males and females were included in each age decade to ensure full coverage of the adult lifespan. All participants gave written informed consent prior to the onset of data collection. The local ethics board approved the study.

### 4.2 MRI scanning

Images were acquired at the Spinoza Centre for Neuroimaging in Amsterdam, the Netherlands, using a Philips Achieva 7 T MRI scanner with a 32-channel phased-array coil. Routine quality checks were performed on a weekly basis at the scanner site. T1-weighted, T2* contrasts were obtained using a MP2RAGEME (multi-echo magnetization-prepared rapid gradient echo) sequence (Caan et al., 2019). The MP2RAGEME is an extension of the MP2RAGE sequence (Marques et al., 2010) and consists of two rapid gradient echo (GRE_1,2_) images that are acquired in the sagittal plane after a 180 degrees inversion pulse and excitation pulses with inversion times TI_1,2_ = [670 ms, 3675.4 ms]. A multi-echo readout was added to the second inversion at four echo times (TE_1_ = 3 ms, TE_2,1-4_ = 3, 11.5, 19, 28.5 ms). Other scan parameters include flip angles FA_1,2_ = [4, 4] degrees; TR_GRE1,2_ = [6.2 ms, 31 ms]; bandwidth = 404.9 MHz; TR_MP2RAGE_ = 6778 ms; acceleration factor SENSE PA = 2; FOV = 205 × 205 × 164 mm; acquired voxel size = 0.7 × 0.7 × 0.7 mm; acquisition matrix was 292 x 290; reconstructed voxel size = 0.64 × 0.64 × 0.7 mm; turbo factor (TFE) = 150 resulting in 176 shots; Total acquisition time = 19.53 min.

### 4.3 Quantitative MRI modeling and parcellation

The MP2RAGEME consists of two interleaved MPRAGEs with different inversions and four echoes in the second inversion. Based on these images, we estimated quantitative MR parameters of R1, R2* and QSM as follows. First, we took advantage of the redundancy in the MP2RAGEME sequence to perform a PCA-based denoising with LCPCA (Bazin et al., 2019). R1 maps were then computed using a look-up table (Marques et al., 2010). R2*-maps were computed by least-squares fitting of the exponential signal decay over the four echoes of the second inversion. QSM images were obtained from the phase maps of the second, third, and fourth echoes of the second inversion with TGV-QSM (Langkammer et al., 2015). Skull stripping, required for QSM, was performed on the second inversion, first echo magnitude image (Bazin et al., 2014).

The anatomical regions of interest were defined with the MASSP automated algorithm (Bazin et al., 2020) on the basis of the R1, R2* and QSM image maps. The algorithm combines location, shape, and quantitative MRI priors to define 17 subcortical anatomical regions, listed in Table 1. Separate masks for left and right hemisphere were obtained except for 3V, 4V, and *fx*.

For this study, the MASSP algorithm was trained on renormalized versions of the quantitative contrasts using a fuzzy C-means clustering of intensities, and linearly interpolating between cluster centroids. The renormalized contrasts were thus less sensitive to the intensity variations induced by aging. Additionally, the registration to the MASSP atlas was performed in two successive steps, producing more accurate alignment of the anatomical priors with each subject.

### 4.4 Iron and myelin approximation

Iron and myelin are main determinants of MR image contrast (Stüber et al., 2014). Several lines of research indicate that the concentrations of iron and myelin are approximately linearly related to qMRI metrics R1, R2* and QSM (Hametner et al., 2018; Mangeat et al., 2015; Marques et al., 2017; Metere & Möller, 2018; Rooney et al., 2007; Stüber et al., 2014). Assuming a linear relationship between iron and myelin on the one hand, and qMRI on the other, linear models can be fit and used to predict iron and myelin contents based on qMRI values (Metere & Möller, 2018):

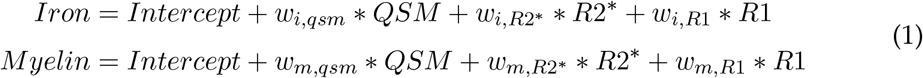

Estimating the parameters *w* of these models requires population-average estimates of iron and myelin content for a variety of regions of interest that cover the range of R1, R2*, and QSM values observed across the brain. Following the approach by Metere and Möller (2018), we obtained these values from the literature (Hallgren & Sourander, 1958; Metere & Möller, 2018; Randall, 1938), and supplemented those values using observations in *post mortem* tissue (detailed below). For iron estimates, Hallgren and Sourander (1958) provided quantifications in regions of interest that vary in both iron contents and qMRI values, which allows for stable estimators of the weights in Equation 1. An iron concentration of approximately 0 in the ventricles was assumed.

As a reference for myelin concentrations, we used work by Randall (1938), which provides lipid concentrations for the corona radiata, frontal and parietal white matter, brain stem, thalamus, caudate, and frontal and parietal grey matter. Following Metere and Möller (2018), we assumed that these lipid concentrations reflect myelin concentrations. Unfortunately, the reported regions do not include iron-rich nuclei, which limits the range of (especially) R2* and QSM values with known corresponding lipid concentrations. Using a limited number of regions of interest to estimate the myelin model could limit the generalizability of the estimated parameters to structures with lower R2* and/or QSM values, which would bias myelin estimates in iron-rich structures like some basal ganglia nodes (e.g., based on using only Randall’s (1938) lipid concentrations, Metere and Möller (2018) obtained *negative* myelin concentrations in various basal ganglia structures).

To supplement the literature-based myelin concentrations, we approximated the myelin contents of other regions of interest using a *post mortem* specimen. Here, we made the following assumptions:

1. The optic density of tissue in our silver stains is approximately linearly related to the concentration of myelin in that tissue in our regions of interest (see Figure 8). Here, we confirmed that silver stains were not saturated even in the white matter regions;
2. The myelin concentrations in the *post mortem* specimen are representative of the population and comparable to the concentrations reported by Randall (1938);
3. The myelin concentrations in white matter reported by Randall (1938) are comparable to the myelin concentration in the internal capsule. Similarly, the myelin concentrations in parietal cortex are comparable to insular cortex.

**Figure 8:**
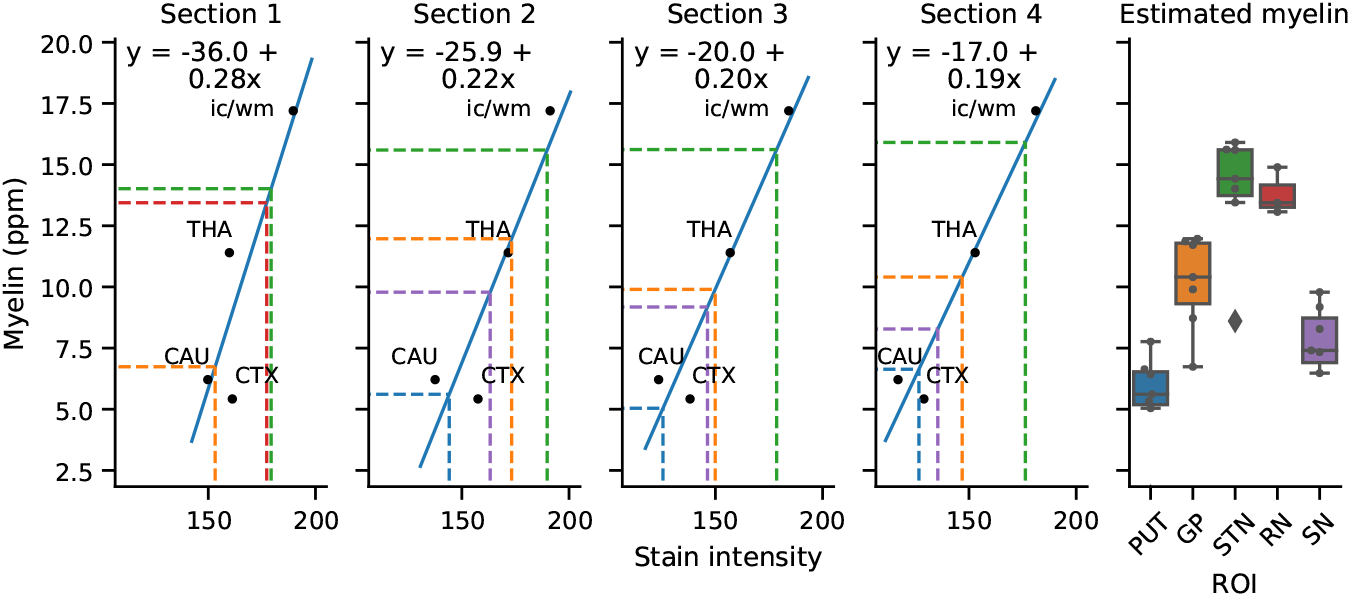
Procedure of estimating myelin contents using a *post mortem* specimen. In each section (seven in total, first four shown), the stain intensities corresponding to CAU, THA, insular cortex (CTX) and the internal capsule (ic/wm) were estimated. For each section individually, a calibration curve was estimated to map stain intensity to myelin values (solid blue lines and equations). Within the range of interest, the relation between stain intensity and myelin content could be approximated with a linear trend. Then, the intensity values for putamen (PUT), GP, STN, RN, and SN were estimated per section (colored dashed lines; note that not all sections contained all structures), and the corresponding myelin values were calculated. Boxplots in the right panel show across-section variability in estimated myelin contents and suggest agreement across sections. The center line in each box marks the median, box limits are the across-section interquartile range, and whiskers are at 1.5 times the interquartile range below and above the box limits. ROI = Region of interest.

Seven 200 *μ*m coronal sections of a single specimen were stained according to the method described by Bielschowsky (for details, see Alkemade, Pine, et al., 2020). Sections included the caudate nucleus, thalamus, internal capsule, and insular cortex, in which we estimated the median intensity of the lightness of the stain (the optic density). Randall (1938) reports quantified lipid concentrations of the caudate nucleus and thalamus, which can be directly compared to the stain intensities, as well as of parietal grey and white matter. The caudate nucleus, thalamus, and parietal grey and white matter (as reported by Randall (1938)) were not visible in the same histological section, and we therefore used insular cortex as a reference region for grey matter, and the internal capsule as a reference region for white matter. For each section separately, we then created a linear calibration curve, which allowed us to determine lipid concentrations based on the stain intensity (Figure 8) for putamen, globus pallidus, subthalamic nucleus, red nucleus, and substantia nigra.

For the region of which population-averaged iron and myelin contents were known, we estimated the qMRI values using the MRI data. Median qMRI values were calculated using the MASSP masks for subcortical regions, and a MGDM and CRUISE parcellation was used to obtain individual masks for brain stem, cerebellum, and cortex (Bazin et al., 2014). We included only participants of 30 years and older to match the ages of the specimens on which the iron and myelin estimates are based. For brain regions for which we had estimated the myelin content using our *post mortem* specimen, we only included AHEAD subjects of 70 years and older (17 participants total) to approximately match ages of the MRI data and the specimen. Tables S1 and S2 list the iron and myelin concentrations, respectively, and their corresponding qMRI values, that were used to estimate the parameters in Equations 1.

To test whether all qMRI metrics were required as predictors to accurately predict iron and myelin content, we fitted linear models with all eight possible combinations of R1, R2*, and QSM. Models were fitted using ordinary least squares (OLS). For each model, we estimated the Akaike information criterion (AIC; Akaike, 1973) to identify the model that is expected to have the highest predictive performance, and used the model with lowest AIC values (AIC and BIC values agreed on the winning model). We used the AIC here instead of the BIC as the AIC is expected to select models with the highest cross-validated predictive performance, whereas the BIC is expected to select the data-generating model (Wagenmakers & Farrell, 2004). The parametrized winning models were as follows:

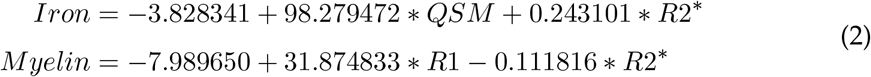

Figure 9 visualizes quality of fit of the winning models and shows that a high proportion of variance in myelin and iron can be explained using the qMRI values. Note that the model weights cannot directly be compared to the weights from Stüber et al. (2014), which were obtained using formalin fixated *post mortem* tissue. Formalin fixation can change qMRI values (Birkl et al., 2016; Langkammer et al., 2012; Schmierer et al., 2008; T. M. Shepherd et al., 2009; Tovi & Ericsson, 1992). A second complicating factor is that qMRI values can vary between MRI sites (Mancini et al., 2020), suggesting the need to re-estimate model weights when using qMRI obtained at a different site.

**Figure 9:**
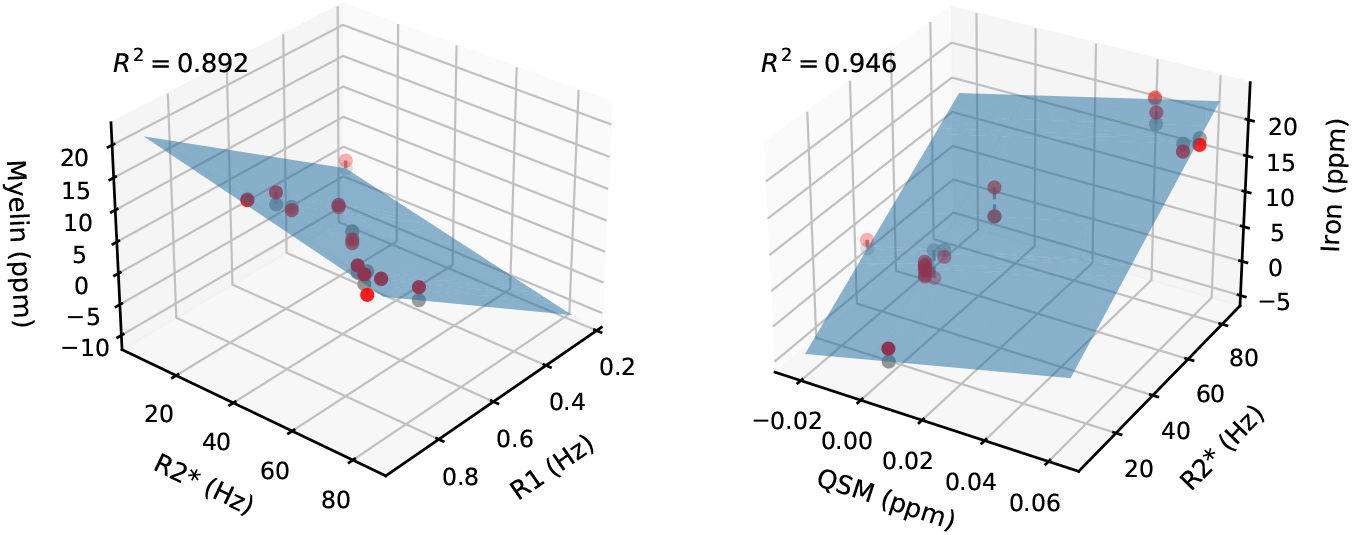
Quality of fit of the myelin (left) and iron (right) model. The planes are given by Equations 2. Red dots illustrate data points, grey dots are the model predictions for these data points.

Using these simplified biophysical models, we calculated whole-brain iron and myelin maps, and obtained participant-specific myelin and iron values for all structures using the MASSP masks. Iron and myelin maps of a representative participant are shown in Figure 10.

**Figure 10:**
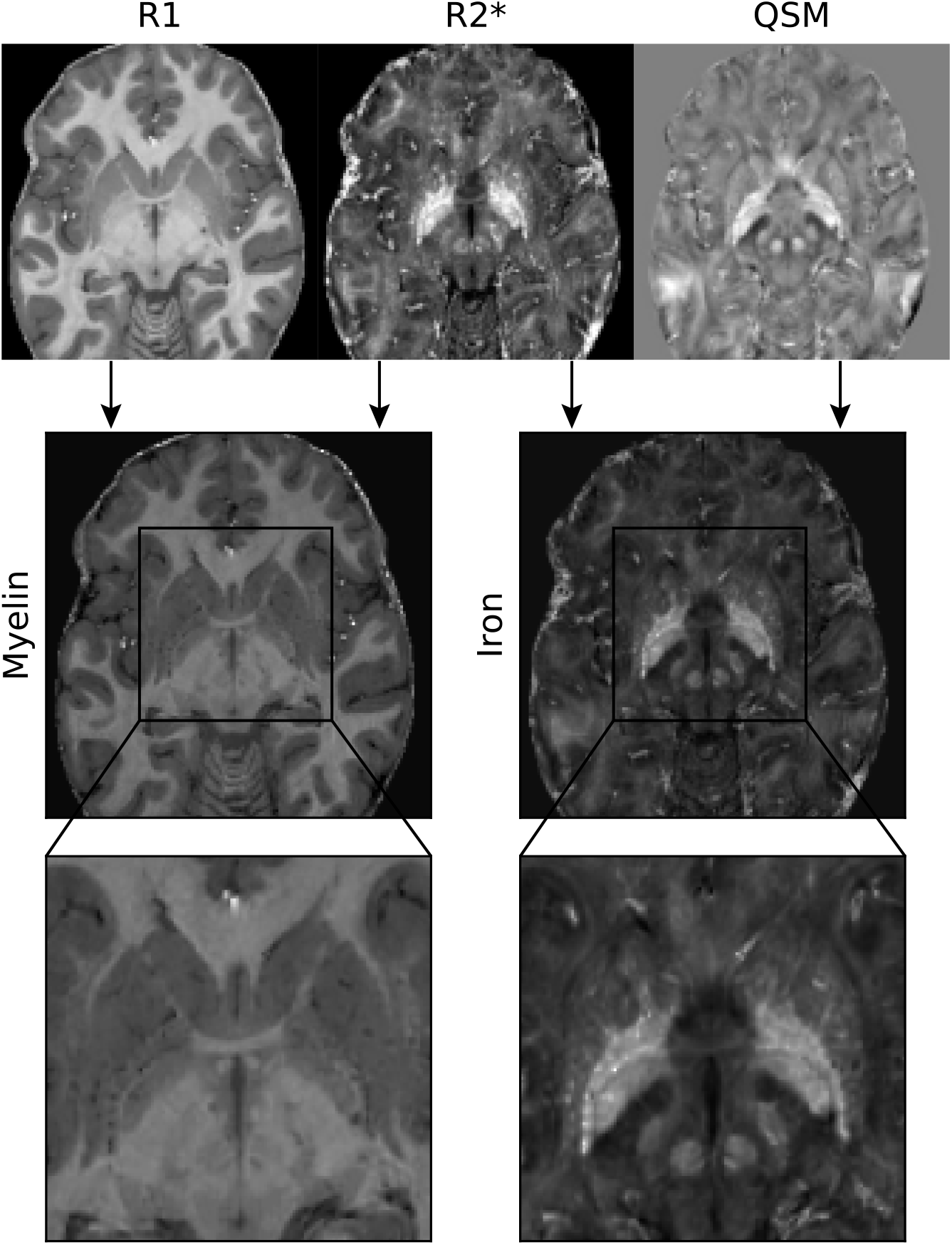
Example of myelin (left) and iron map (right) of a representative participant. The top row shows the R1, R2*, and QSM maps, which were linearly combined into myelin and iron maps (middle and bottom row) using Equations 2. Note the hyperintense appearance of iron rich structures such as the rounded shape of the red nucleus.

It is important to emphasize that the iron and myelin estimates we report are based on simplifying assumptions with regard to the linearity of the relation between qMRI and iron/myelin, and on the iron/myelin concentrations on which the biophysical models are fitted (detailed above). As such, the iron and myelin estimates should be not be interpreted as absolute measurements, but rather as approximations that serve to guide the interpretations of qMRI values in terms of the most likely underlying biological contributors to those values.

### 4.5 Thickness estimation

We calculated local structure thickness based on a medial skeleton representation: for each structure, we estimated the skeleton as the ridge equidistant to the structure boundaries. Thickness was defined as twice the distance between the skeleton and the closest boundary, using the method described in Bazin et al. (2020). In other words, local thickness measures at every location inside the structure the distance between the two closest boundaries of that structure, extending the concept of cortical thickness to more complex shapes. Contrary to volume, thickness can be determined at the position of each voxel within a structure, thus providing local information. A similar thickness measure was also used in Ho et al. (2020) to detect subtle shape differences.

### 4.6 Center of mass

For all structures, we calculated the center of masses in Cartesian x, y, and z coordinates per participant after an affine transformation to group space, using ANTs (Avants, Epstein, Grossman, & Gee, 2008) with mutual information. The affine transformation was applied to correct for inter-individual differences in intracranial volume and neurocranium shape, while retaining inter-individual variability in anatomy relative to the neurocranium.

### 4.7 Age-related change modelling

We describe the age-related changes in iron concentration, myelin content, volume, and thickness, as well as in the center of mass in x, y and z coordinates. For iron, myelin and thickness, we report both a median reflecting the central tendency and interquartile range reflecting structure homogeneity. For thickness, the interquartile range reflects the within-structure variability of thickness, quantifying the regularity of the shape. We also analyzed the R1, R2*, and QSM values, which can be found in the online app (https://subcortex.eu/app).

Exploratory modelling of the between-hemisphere differences per structure suggested no between-hemisphere difference in aging patterns for most structures. Therefore, we subsequently assumed that the age-related changes in each structure were the same in both hemispheres, to reduce the total number of models fitted. We collapsed across hemispheres by taking the mean value across both hemispheres per structure and participant.

Prior to fitting the aging models, we excluded outliers based on their Mahalanobis distance (cut-off 10.827, corresponding to *p* < 0.001, 0.69% of all data points). Per ROI and dependent variable, we then fit the following set of 24 potential models, with all possible combinations of the following predictors: A linear influence of age, a quadratic influence of age, sex, an interaction between age and sex, and an interaction between a quadratic influence of age and sex. We excluded models with both interaction terms, as this would imply implausibly large between-sex differences in aging patterns.

Models were fit with OLS as implemented in statsmodels (Seabold & Perktold, 2010) for the Python programming language. Models were compared with the Bayesian Information Criterion (BIC; Schwarz, 1978), which quantifies the quality of fit penalized for model complexity. Lower BIC values indicate more parsimonious trade-offs between quality of fit and model complexity and are preferred. Based on the winning model, we removed influential data points using Cook’s distance (cut-off 0.2, 0.18% of all data points; we used a more conservative cut-off than 4/*n*, which is sometimes recommended [Rawlings, Pantula, and Dickey, 1998]). We then refitted all models on the data excluding the influential data points, and performed a new model comparison.

Using the winning age-related change models, we quantified the total age-related changes by summing the yearly absolute changes between 19 and 75 years old (Figures 2–4, S2), divided by the model’s prediction at 19 years old. This quantity summarizes the amount of change relative to the baseline value at 19, and allows for comparing models with different specifications (e.g., linear vs quadratic models). For winning models that included sex as a predictor (or interactions between sex and age), the cumulative yearly absolute changes were calculated for both sexes and then averaged. For models where the overall direction of the age-related change was negative (identified by comparing the predicted values at 19 and 75 years old), we took the negative of the total age-related changes. As our data contains only a single data point above 75 years old, we chose to limit the range of ages that we interpret to 19–75 years old, where we expect our models’ predictions to be most robust.

To identify similarities in the aging patterns of groups of regions, we calculated all pairwise correlations between regions (based on Figure S2 as input, excluding the location changes)). The resulting correlation matrix was ordered using hierarchical clustering with the UPGMA algorithm with the Euclidean distance as a metric (as default settings in the seaborn visualization package, Waskom and development team, 2020), shown in Figure 7.

### 4.8 Confidence intervals and standard errors

Confidence intervals in Figure 1 were obtained using a bootstrapping procedure with 10,000 iterations. We iteratively sampled 105 random observations with replacement from the data, based on which we estimated the median, and took the 2.5^th^ and 97.5^th^ percentile of the 10,000 medians as the 95% confidence interval. The standard errors in Figure 6 were obtained using a similar bootstrapping procedure, in which winning model specifications were iteratively fit on 10,000 random samples (drawn with replacement) from the data. Per iteration, cumulative change metrics were estimated. The standard deviation of the cumulative change metrics across iterations was used as an estimator of the standard error. For winning models that do not include age as a predictor variable, the standard error is 0 since the cumulative age-related change metric is 0 in each iteration.

## Open Science

A prior version of the individual qMRI maps has been released as AHEAD (Alkemade, Mulder, et al., 2020) can be found at https://doi.org/10.21942/uva.10007840.v1. All derived participant-wise and region-wise measures can be downloaded from our app at https://subcortex.eu/app. All code used to estimate the models and produce the figures can be found at https://osf.io/mvdbe/.

## Acknowledgements

This research was financially supported by STW/NWO (#14017; BUF and AA), ERC PoC (BUF), NWO Vici (016.Vici.185.052; BUF).

## CRediT author statement

**Steven Miletić**: Conceptualisation, Methodology, Software, Validation, Formal analysis, Investigation, Data curation, Writing – Original draft, Writing – Review & Editing, Visualisation, **Pierre-Louis Bazin**: Conceptualisation, Methodology, Investigation, Software, Supervision, Writing – Original draft, Writing –Review & Editing, Formal analysis, Data curation, **Scott Isherwood**: Writing – Review & Editing, **Max C. Keuken**: Writing – Review & Editing, **Anneke Alkemade**: Conceptualisation, Methodology, Validation, Investigation, Resources, Data curation, Writing – Original draft, Writing – Review & Editing, Supervision, Project administration, Funding acquisition, **Birte U. Forstmann**: Conceptualisation, Methodology, Validation, Investigation, Resources, Data curation, Writing – Review & Editing, Supervision, Project administration, Funding acquisition.

## Supplementary Materials

### CNR comparisons

Under the assumption of mono-exponential signal decay, the expected contrast-to-noise ratio (CNR) of an T2*-weighted single echo, echo planar imaging protocol, per unit change in T2*, is given by (e.g. Poser, Versluis, Hoogduin, & Norris, 2006; Posse et al., 1999):

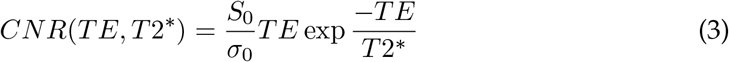

where *S*_0_ and *σ*_0_ are the signal and (temporal) variance, respectively, at echo time *TE* = 0. In the following comparisons, we assume that *S*_0_ and *σ*_0_ are the same across structures and protocols. This assumption is unlikely to be true for between-region CNR comparisons, as *S*_0_/*σ*_0_ is typically lower in subcortical regions compared to cortical regions in part due to the larger distance to the MRI receiver coils. We ignore any potential between-region differences in the size of the T2* changes that result from changes in the oxygenation levels. Under these assumptions, CNR ratios can be used to compare the expected CNRs when using echo time *A* to study region *n* with the expected CNR when using echo time *B* to study region *m*:

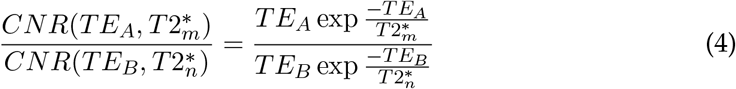

First, we can compare the expected CNR in the red nucleus with the CNR in the amygdala, both in 19 years old participants, using T2* values obtained through our app (https://subcortex.eu/app). Filling in 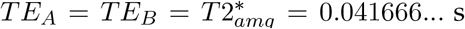 (i.e., in both protocols, we use an echo time optimised for the amygdala at 19 years old), and 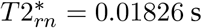, we find 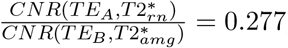, implying the CNR in the red nucleus is approximately 72% lower than in the amygdala with this echo time. In practice, since *S*_0_/*σ*_0_ is also likely to be lower in the red nucleus than in the amygdala, the CNR ratio will be even lower.

Second, we can estimate the effect of age-related decreases in T2*. The red nucleus has an approximate *T* 2* = 0.018 s at 19 years old, and *T* 2* = 0.0137 s at 50 years old. If an echo time of *TE_A_ = TE_B_* = 0.018 s is used for a participant of 50 years old, then 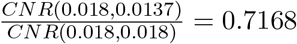, showing an approximately 28% loss in CNR compared to the CNR that would be obtained with this echo time in a participant of 19 years old.

Third, we can compare the expected CNR in the red nucleus at 19 years old (*T* 2* = 0.018 s) when using the optimal echo time *T E_A_* = 0.018 s with the CNR that would be obtained if an echo time is used that is optimal for the amygdala: *T E_B_* = 0.0042 s. The ratio 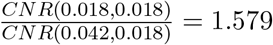, showing that a substantial CNR gain can be expected from optimising the echo time to meet the specific requirements of that region.

Finally, we can compare the effect of adapting echo times to adjust for age-related changes in T2*. Taking again the red nucleus as an example, the T2* decreases from 0.018 s to 0.0138 s between 19 and 50 years old. Suppose we analyze the red nucleus in a 50 year old participant using the corresponding optimal echo time (hence, *T E_A_* = *T* 2* = 0.0137 s), then, compared with using an echo time optimal for young participants *T E_B_* = 0.018, we find that 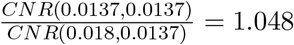. Hence, adjusting for the age-related decrease in T2* leads to modest CNR gains.

In practice, changing the echo time may not always be possible due to hardware limitations (e.g., the slew rate of the MRI gradients limits the minimum echo time that can be achieved), and potentially requires undersampling of k-space (e.g., using GRAPPA, SENSE, or partial Fourier) or bandwidth changes. These additional changes will affect the protocols’ *S*_0_/*σ*_0_, complicating direct comparison between the expected performance of two candidate MRI protocols. Another option is to use multi echo protocols, in which data is acquired at multiple echo times, which can be optimised for multiple regions at the same time (Gowland & Bowtell, 2007; Kundu et al., 2017; Miletić, Boag, & Forstmann, 2020; Puckett et al., 2018). These additional factors should be taken into account when developing an MRI protocol in order to find the optimal trade-off between echo time settings and *S*_0_/*σ*_0_.

**Figure S1:**
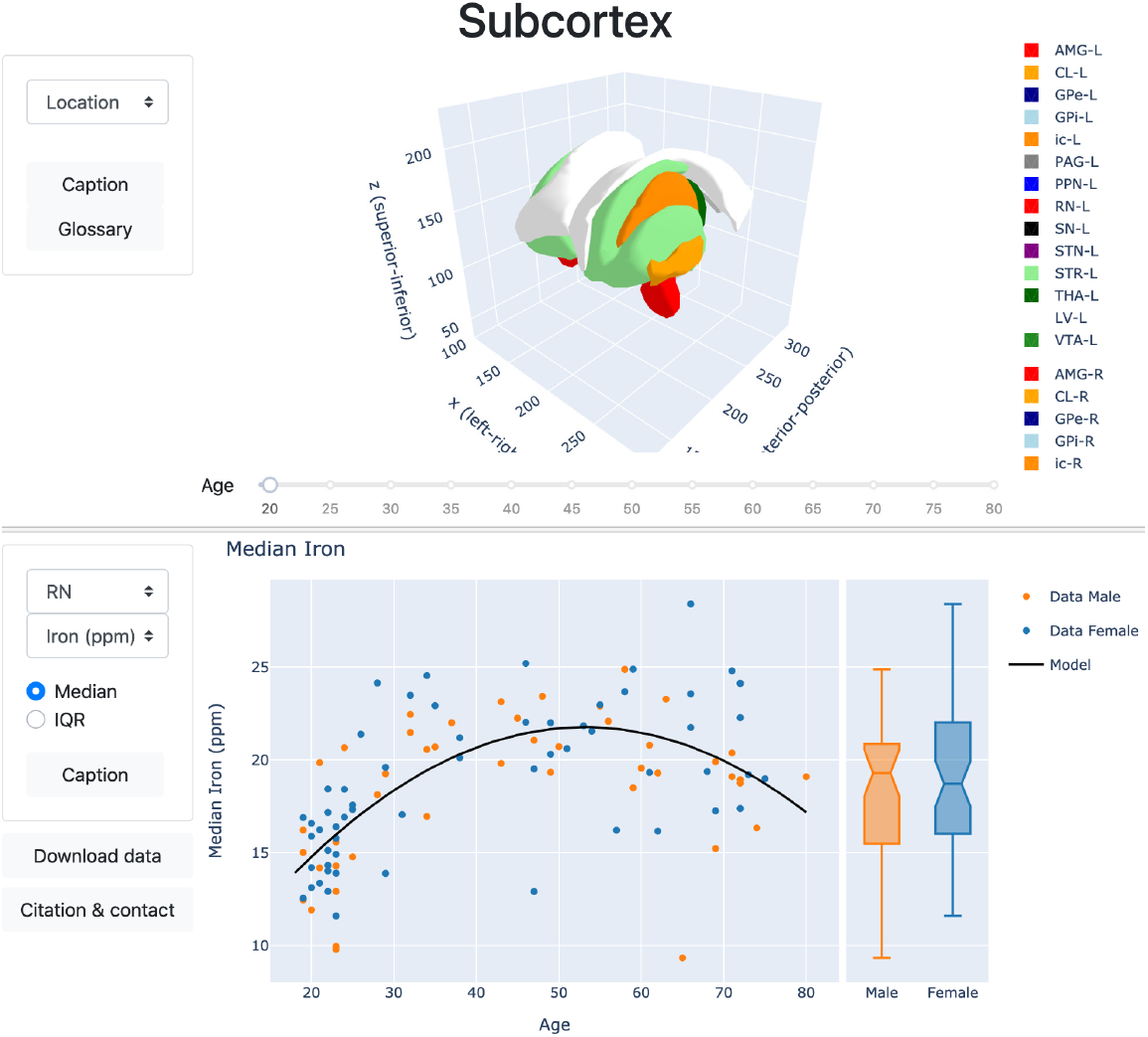
Illustration of the interactive app which allows for visualizing all data and agerelated change models. Top panel features a 3D mesh plot that includes all 17 subcortical structures under investigation. The age slider can be used to visualize the age-related changes in center of mass location (currently visible), or the median or interquartile range of iron, myelin, R1, R2*, and QSM values, color-coded on the structures. Bottom panel features scatterplots of the relation between age and twelve measures (median and interquartile range of myelin, iron, R1, R2*, QSM, and thickness) for all structures. Optional captions are included, and the data underlying the bottom panel can readily be downloaded as a csv file. The app can be accessed via https://subcortex.eu/app

**Table S1:**
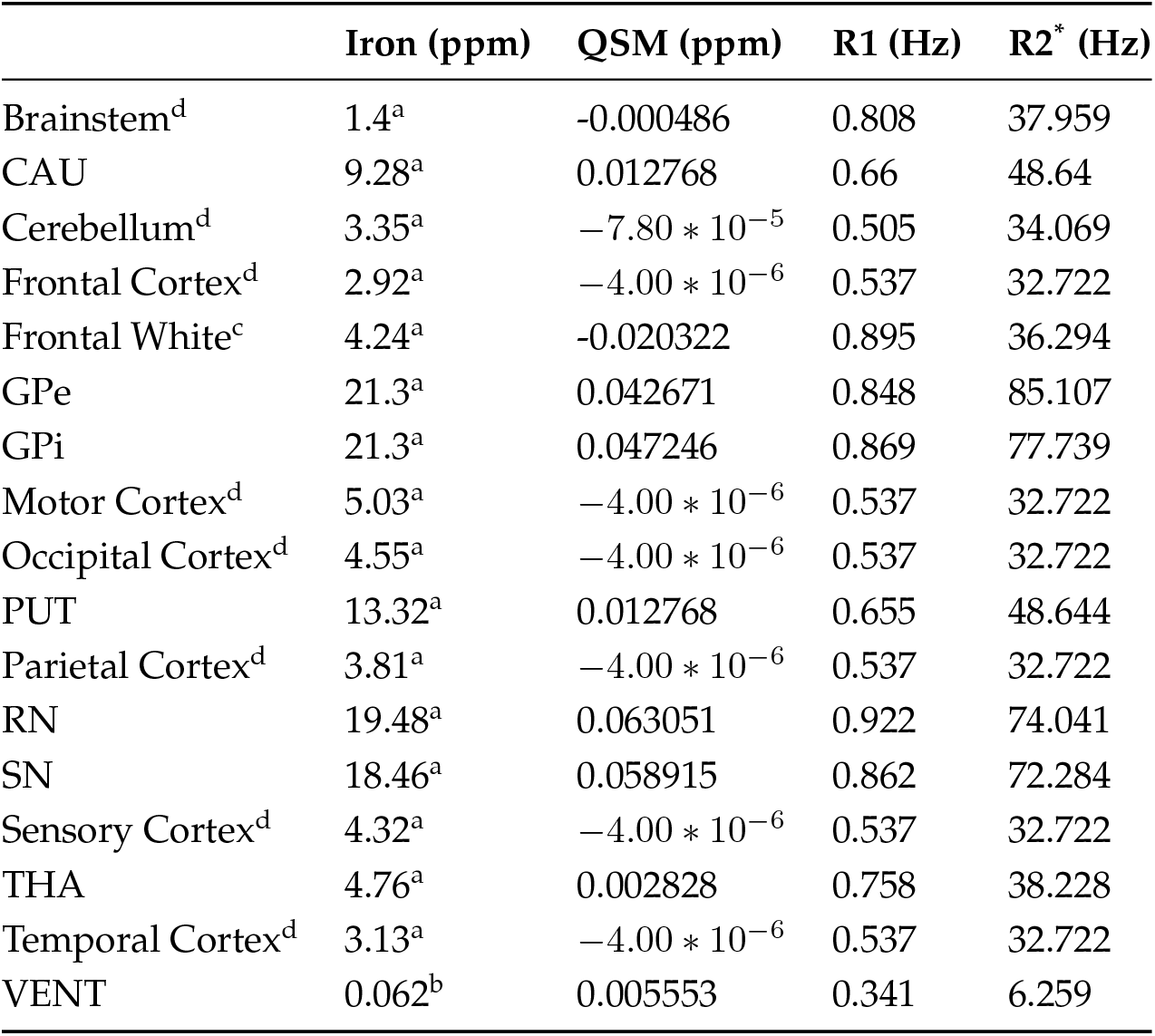
Iron estimates and corresponding qMRI values used for estimating linear model Equation 1. For all regions that were divided in subregions in one but not the other source (i.e. GP vs GPi/GPe; STR vs PUT/CAU; brain stem vs medulla oblongata; and cortical areas), we entered all subregions in the OLS model, using the more specific values where possible, and the global values otherwise. E.g., both GPe and GPi were in the model and shared iron values (Hallgren and Sourander report only GP), but different qMRI values. ^a^ From Hallgren and Sourander (1958); ^b^ cerebrospinal fluid estimates from Metere and Möller (2018); ^c^ We used the qMRI metrics of the internal capsule here; ^d^ Cortex, brain stem, cerebellum were parcellated using MGDM and CRUISE Bazin et al. (2014).

|  | Iron (ppm) | QSM (ppm) | R1 (Hz) | R2* (Hz) |
| --- | --- | --- | --- | --- |
| Brainstem<sup>d</sup> | 1.4<sup>a</sup> | -0.000486 | 0.808 | 37.959 |
| CAU | 9.28<sup>a</sup> | 0.012768 | 0.66 | 48.64 |
| Cerebellum<sup>d</sup> | 3.35<sup>a</sup> | $-7.80 * 10^{-5}$ | 0.505 | 34.069 |
| Frontal Cortex<sup>d</sup> | 2.92<sup>a</sup> | $-4.00 * 10^{-6}$ | 0.537 | 32.722 |
| Frontal White<sup>c</sup> | 4.24<sup>a</sup> | -0.020322 | 0.895 | 36.294 |
| GPe | 21.3<sup>a</sup> | 0.042671 | 0.848 | 85.107 |
| GPi | 21.3<sup>a</sup> | 0.047246 | 0.869 | 77.739 |
| Motor Cortex<sup>d</sup> | 5.03<sup>a</sup> | $-4.00 * 10^{-6}$ | 0.537 | 32.722 |
| Occipital Cortex<sup>d</sup> | 4.55<sup>a</sup> | $-4.00 * 10^{-6}$ | 0.537 | 32.722 |
| PUT | 13.32<sup>a</sup> | 0.012768 | 0.655 | 48.644 |
| Parietal Cortex<sup>d</sup> | 3.81<sup>a</sup> | $-4.00 * 10^{-6}$ | 0.537 | 32.722 |
| RN | 19.48<sup>a</sup> | 0.063051 | 0.922 | 74.041 |
| SN | 18.46<sup>a</sup> | 0.058915 | 0.862 | 72.284 |
| Sensory Cortex<sup>d</sup> | 4.32<sup>a</sup> | $-4.00 * 10^{-6}$ | 0.537 | 32.722 |
| THA | 4.76<sup>a</sup> | 0.002828 | 0.758 | 38.228 |
| Temporal Cortex<sup>d</sup> | 3.13<sup>a</sup> | $-4.00 * 10^{-6}$ | 0.537 | 32.722 |
| VENT | 0.062<sup>b</sup> | 0.005553 | 0.341 | 6.259 |

**Table S2:** For all regions that were divided in subregions in one but not the other source (i.e. GP vs GPi/GPe; STR vs PUT/CAU; brain stem vs medulla oblongata; and cortical areas), we entered all subregions in the OLS model, using the more specific values where possible, and the global values otherwise. E.g., both CAU and PUT were in the model with separate myelin values, but identical qMRI values since the MASSP parcellation only reports STR. ^a^ From Hallgren and Sourander (1958); ^b^ cerebrospinal fluid estimates from Metere and Möller (2018); ^c^ We used the qMRI metrics of the internal capsule here; ^d^ Cortex, brain stem, cerebellum were parcellated using MGDM and CRUISE Bazin et al. (2014).

|  | Myelin (ppm) | QSM (ppm) | R1 (Hz) | R2* (Hz) |
| --- | --- | --- | --- | --- |
| Brainstem<sup>d</sup> | 15.36<sup>a</sup> | -0.000486 | 0.808 | 37.959 |
| CAU | 6.21<sup>a</sup> | 0.012768 | 0.655 | 48.644 |
| Frontal Cortex<sup>d</sup> | 5.08<sup>a</sup> | $-4.00 * 10^{-6}$ | 0.537 | 32.722 |
| Frontal White<sup>c</sup> | 16.26<sup>a</sup> | -0.020322 | 0.895 | 36.294 |
| GPe | 10.404<sup>a</sup> | 0.044293 | 0.813 | 84.875 |
| GPi | 10.404<sup>a</sup> | 0.046384 | 0.838 | 75.216 |
| Parietal Cortex<sup>d</sup> | 5.42<sup>a</sup> | $-4.00 * 10^{-6}$ | 0.537 | 32.722 |
| PUT | 5.611<sup>a</sup> | 0.012768 | 0.655 | 48.644 |
| RN | 13.442<sup>a</sup> | 0.059001 | 0.894 | 72.979 |
| SN | 7.404<sup>a</sup> | 0.05624 | 0.846 | 71.532 |
| STN | 14.423<sup>a</sup> | 0.058185 | 0.935 | 78.924 |
| THA | 11.4<sup>a</sup> | 0.002828 | 0.758 | 38.228 |
| VENT | 0.002<sup>b</sup> | 0.005553 | 0.232 | 6.259 |

**Figure S2:**
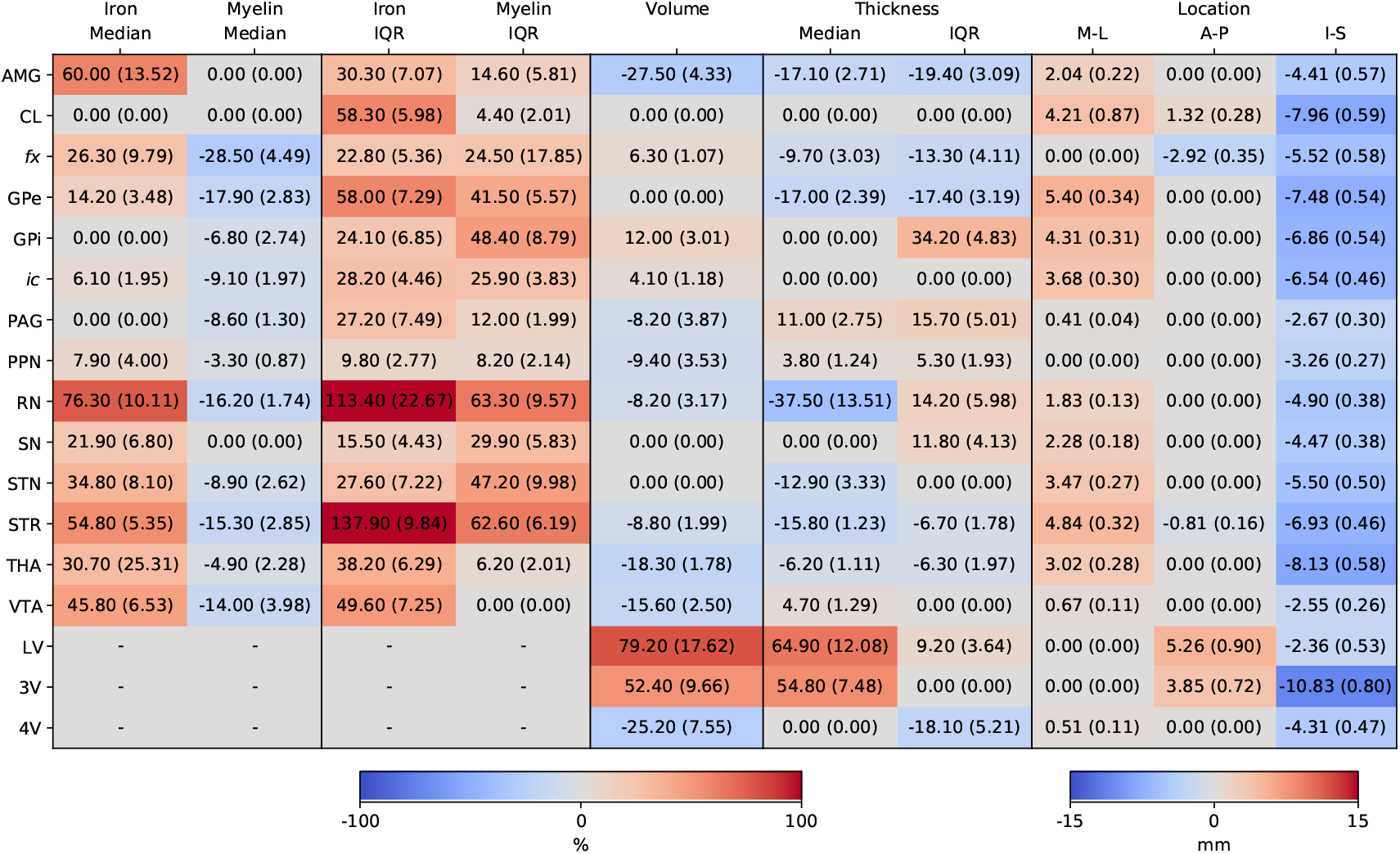
Matrix of cumulative change in all dependent variables (columns) for individual structures (rows). Values indicate the summed absolute change between 19-75 years old. Negative values indicate decreases. Standard errors were obtained by bootstrapping with 10,000 iterations and are shown in parentheses. Location change in fornix was not included since the quality of parcellation of the fornix posterior horns increases with age, resulting in artificial shifts in the estimated center of mass with age. The ventricular system is assumed to have no iron or myelin concentration and is excluded from analysis.

